# An optimised method for the recovery of laboratory generated SARS-CoV-2 aerosols by plaque assay

**DOI:** 10.1101/2022.10.31.514483

**Authors:** Rachel L Byrne, Susan Gould, Thomas Edwards, Dominic Wooding, Barry Atkinson, Ginny Moore, Kieran Collings, Cedric Boisdon, Simon Maher, Giancarlo Biagini, Emily R Adams, Tom Fletcher, Shaun H Pennington

**Affiliations:** Centre for Drugs and Diagnostics, Department of Tropical Disease Biology, Liverpool School of Tropical Medicine, Liverpool, UK; UK Health Security Agency, Porton Down, UK; Department of Electrical Engineering and Electronics, University of Liverpool, Liverpool, UK

## Abstract

We present an optimised method for the recovery of laboratory generated SARS-CoV-2 virus by plaque assay. This method allows easy incorporation into existing standard operating procedures of biological containment level 3 (BCL3) laboratories.

## Introduction

The perceived risk of the aerosolisation of SARS-CoV-2 has been debated since its emergence in December 2019 [1]. Whilst convincing, the argument for aerosol transmission of COVID relies almost exclusively on retrospective analysis of outbreaks or superspreading events [2], with a notable absence of laboratory confirmed, quantifiable, viral isolation data. Plaque assays remain the gold standard method for viral quantification through the enumeration of discrete plaques in cell culture which can be counted to infer viral titre of the inoculum [3]. While cytopathic assays also demonstrate the presence of infectious material, they are subjective, and rely on operator expertise to accurately detect changes in cell morphology caused by infecting virus [4]. We present an optimised method for the recovery of laboratory generated SARS-CoV-2 aerosols utilising gelatine membranes by plaque assay.

Establishing a reliable method for capturing SARS-CoV-2 from the air remains a substantial experimental challenge. A small number of studies have detected viral RNA from the air of hospital rooms [5–8], with only one group able to demonstrate recovery of viable virus [9,10]; however, to date, recovery and quantification of aerosolised SARS-CoV-2 has yet to be demonstrated. Here, we utilise a portable air sampler that is battery powered and has proven effective at capturing SARS-CoV-2 RNA from hospital rooms [11]. Whilst viral recovery optimisation has been demonstrated by the manufacturer (Sartorius, Germany) the method relies on mechanical agitation of the membrane and the additional of chemicals. This may not be appropriate in biological containment level 3 (BCL3) laboratories due to the risk of aerosol generation and/or spills, especially if working in an open-fronted, class II microbiological safety cabinet (MBSC) [12].

The main costs associated with plaque assays, excluding BCL3 laboratory maintenance, is the upkeep of cell lines for viral inoculation and plate production which is both cost- and labour-intensive. Thus, cell culture is typically reserved for clinical samples. Therefore, an efficient method for the analysis of environmental surveillance samples is needed to further our understanding of environmental transmission of SARS-CoV-2 as well as other novel and re-emerging infectious diseases.

## Methods

### Aerosolisation and recovery of SARS-CoV-2

A stock concentration of 1.4 x 10^5^ pfu/mL SARS-CoV-2/human/GBR/Liv_273/2021 virus of the Delta lineage (Pango B.1.617.2 [13]) was aerosolised in a BLAM atomizer (CH technologies, USA) within a class III MBSC. For each condition and replicate, SARS-CoV-2 aerosols were generated for 4 minutes by running the atomizer at a rate of 18 L/min. Recovery was performed by running the MD8 Airport with gelatine membranes (Sartorius, Germany) at a rate of 30 L/min for a total of 50 L. The materials module was detached from the BLAM and MD8 Airport. The gelatine membrane was then transferred in to a 50 mL Falcon tube containing 10 mL of Dulbecco’s modified Eagle’s medium (DMEM) with 10% fetal bovine solution (FBS; both ThermoFisher, USA) and 50 units/mL of penicillin/streptomycin (Gibco, US). For each variable, 3 biological replicates were performed.

### Optimisation for recovery and culture of SARS-CoV-2 virus

During development of this protocol numerous variables were considered and tested (Figure 1). In ‘Phase 1’, we determined if passage in cells was required prior to plaquing. We also established the optimum duration of time (1 hour, 4 hours or 24 hours) to dissolve gelatine membranes. Finally, we investigated the temporary storage conditions of dissolved membranes in DMEM (room temperature (RT), 4 ^o^C or –20 ^o^C) for each sample.

**Figure 1.**
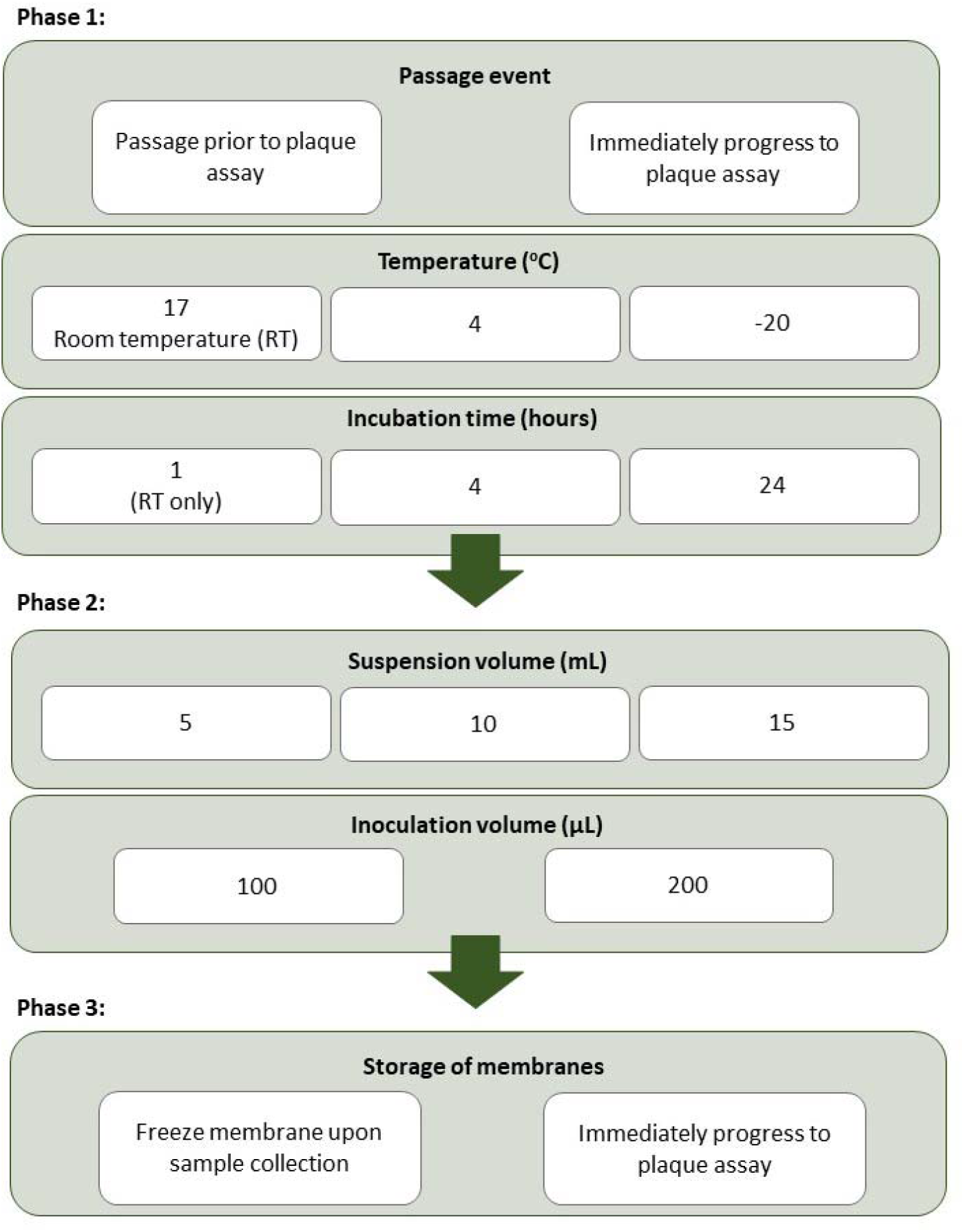
Experimental design for the optimisation of plaque assays for aerosolised SARS-CoV-2.

A passage in cells prior to plaque assays acts an enrichment step but prevents direct quantification from a sample. The storage temperature was investigated as the main variable governing the viscosity of the suspended gelatine membranes, which affects the ability to accurately pipette the suspension. The optimised method was determined based on plaque forming units (PFU) and cycle threshold (Ct) values. In ‘Phase 2’, we focused on the dilution effects of suspension and cell inoculation volumes. This began with the amount of DMEM (5 mL, 10 mL or 20 mL) used to suspend the gelatine membrane following aerosol capture. We also considered the volume of sample used to infect cells (100 μL or 200 μL).

Finally, in ‘Phase 3’, we measured the impact of freezing gelatine membranes immediately after collection to allow for samples to be processed when most convenient for laboratory staff. The full experimental design is shown in Figure 1.

### Quantification of virus

Plaque assays were used for the quantification of virus utilising Vero E6 cells (C1008; African green monkey kidney cells) [14]. 20μL of dissolved membrane/DMEM suspension and 180μl 2% DMEM were added to the first column of a 24-well plate, producing 4 technical replicates. A 1:10 serial dilution was performed with 2% DMEM, and a positive and negative control included in each plate. Plates were incubated at 37 °C for 1 hour and then 1 mL of media overlay (equal parts 2.2% Cellulose solution (Sigma-Aldrich, USA) and 2% DMEM) added. Plates were then incubated at 37 °C for 3 days. Plates were then removed from the incubation, fixed with 100% formaldehyde (Sigma-Aldrich, USA), for one hour, and stained with 1ml/well of 0.25% crystal violet solution (ThermoScientific, USA). After staining for 1 minute, plates were gently washed with water, then air dried for >3 hours. Viral plaques were then visually counted for each well.

Data handling, analysis and statistical comparisons were all performed using R (3.5.5) (R, 2020). Statistical analyses for PFU/mL yield were performed using Kruskal-Wallis nonparametric test with Dunn’s post-hoc test to identify differences in yields using each variable.

### Passage of virus in cell culture

A full 24 well plate, seeded with cells, was used for passaging of samples. A total volume of 200 μL of dissolved membrane/DMEM suspension was added to each well. Plates were incubated at 37 °C for 2 hours and then 200μl of 10% DMEM added. Plates were incubated at 37 °C for 3 days. After incubation, 200 μL from each well (24 per sample) was combined (4.8 mL in total) and transferred to sterile Eppendorf tubes.

## Results

An optimised method for the recovery of laboratory generated SARS-CoV-2 aerosols from gelatin membranes is shown in Box 1 with each variable based on PFU/mL (Table 1).

### Box 1 An optimised method for the recovery of laboratory generated SARS-CoV-2 aerosols from gelatin membranes.

1. Immediately suspend the gelatin membrane in 20mL Dulbecco’s modified Eagle’s medium (DMEM) with 10% fetal bovine solution (FBS; both ThermoFisher, USA) and 50 units/mL of peniciltln/streptornvcin (Glbco, US).
2. Incubate at room temperature for 4 hours.
3. Add 200μL of suspended membrane/DMEM to a 24 well plate seeded with Vero E6 cells.
4. Incubate plates for 2 hours at 37°C.
5. Add 200μl of 10% DMEM.
6. Incubate plates at 37°C for 3 days
7. Add 200μl of supernatant for step 6 and 180μl 2% DMEM to a new 24-well plate seeded with Vero E6 cells.
8. Perform a 1:10 serial dilution with 2% DMEM across the plate.
9. Incubate at 37°C for 1 hour
10. Add 1 mL of media over lay (equal parts 2.2% Cellulose solution (Sigma-Aldrich, USA) and 2% DMEM).
11. Incubate plates at 37°C for 3 days.
12. Fix with 100% formalde hyde for 1 hour.
13. Stain with 1ml/well of 0.25% crystal violet solution.
14. Gently wash with water and air dried for >3 hours.
15. Visually count viral plaques.

**Table 1:** The effect of a cell passage, temperature and suspension time, suspension and inoculation volume and freezing of gelatin membranes on PFU/mL recovery of aerosolised SARS-CoV-2.

| **Phase 1** |  |  |  |  |  |
| --- | --- | --- | --- | --- | --- |
| 3-day passage prior to plaque assay |  | Biological replicates (PFU/mL) |  |  | Mean (PFU/mL) |
|  |  | 1 | 2 | 3 |  |
| Yes |  | 4.63E+06 | 1.88E+06 | 2.75E+06 | 3.08E+06 |
| No |  | 0 | 0 | 0 | 0 |
| *Temperature and suspension time* |  |  |  |  |  |
| Temperature (°C) | Time (hours) | Biological replicates (PFU/mL) |  |  | Mean (PFU/mL) |
|  |  | 1 | 2 | 3 |  |
| 17 | 1 | 4.63E+06 | 1.88E+06 | 2.75E+06 | 3.08E+06 |
|  | 4 | 3.00E+06 | 1.28E+07 | 1.51E+07 | 1.03E+07 |
|  | 24 | 0 | 0 | 0 | 0 |
| 4 | 4 | 0 | 0 | 0 | 0 |
|  | 24 | 0 | 0 | 0 | 0 |
| -20 | 4 | 0 | 0 | 0 | 0 |
|  | 24 | 0 | 0 | 0 | 0 |
| **Phase 2** |  |  |  |  |  |
| Suspension volume (mL) | Inoculation volume (µL) | Biological replicates (PFU/mL) |  |  | Mean (PFU/mL) |
|  |  | 1 | 2 | 3 |  |
| 5 | 100 |  |  |  |  |
|  | 200 |  |  |  |  |
| 10 | 100 | >1.0E+08 | >1.0E+08 | >1.0E+08 | >1.0E+08 |
|  | 200 | >1.0E+08 | >1.0E+08 | >1.0E+08 | >1.0E+08 |
| 20 | 100 | >1.0E+08 | >1.0E+08 | >1.0E+08 | >1.0E+08 |
|  | 200 | >1.0E+08 | >1.0E+08 | >1.0E+08 | >1.0E+08 |
| **Phase 3** |  |  |  |  |  |
| Storage at -80 prior to processing |  | Biological replicates (PFU/mL) |  |  | Mean (PFU/mL) |
|  |  | 1 | 2 | 3 |  |
| Yes |  | 1.56E+07 | 7.13E+06 | 9.63E+06 | 1.08E+07 |
| No |  | >1.0E+08 | >1.0E+08 | >1.0E+08 | >1.0E+08 |

### Phase 1

The inoculum volume applied to the 24 well plate, did not generate any SARS-CoV-2 viral plaques from membranes progressed immediately to quantification (Table 1). Plaques were however visible after a single passage (Figure 2). Thus, a 3-day passage was included for all other variables.

**Figure 2.**
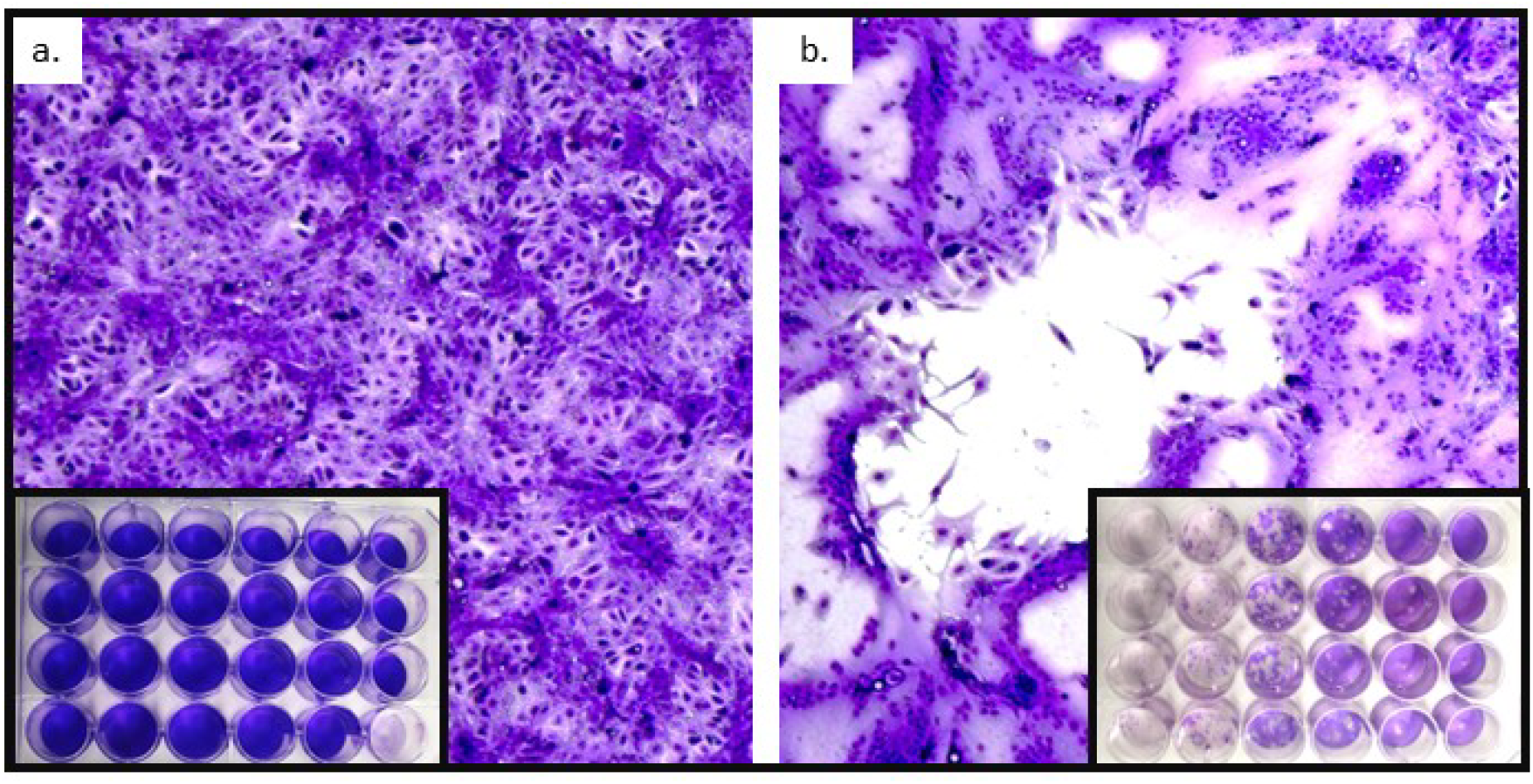
Vero E6 cells infected with aerosolised SARS-CoV-2 stained with crystal violet. **(A**) Representative image of a negative plaque assay – an intact monolayer of Vero E6 cells at 100 × magnification. (B) Representative image of a SARS-CoV-2 plaque - a positive plaque assay of Vero E6 cells at 100 × magnification.

For all temperature and storage time conditions, plaques were only recorded after incubation at RT for 1 hour and 4 hours. Whilst statistically similar (p<0.05) incubation in RT for 4 hours produced the greatest PFU/mL and was used for Phase 2.

### Phase 2

Membranes suspended in 5 mL of DMEM immediately solidified and it was not possible to pipette sufficient volume to inoculate cells and thus samples were discarded. The volume of suspension (10 mL and 20 mL) and inoculation (100 μL and 200 μL) produced similar PFU/mL. As environmental samples are not easily replicated, 20 mL of suspension volume was selected to maximise the amount of sample available for analysis. To mitigate the dilution effect of an increased suspension 200 μL of inoculation volume was preferred.

### Phase 3

Storage of membranes at –80 °C did not have a significant effect on viral recovery (p<0.05) but PFU/mL were lower compared to samples that were immediately processed.

## Discussion

We present an optimised method for the recovery of laboratory generated aerosolised SARS-CoV-2 virus using gelatin membranes by plaque assay. We demonstrate that a single passage in cells enhances recovery, and further demonstrate that the freezing of membranes prior to suspension in culture media reduces viral recovery. Based on the data shown here, we recommend samples are processed immediately after collection. Using a small fraction of volume recovered, we were able to demonstrate successful viral recovery with the inclusion of cell passage prior to plaque assay. Unfortunately, due to the requirement for cell passage, it is not possible to directly quantify viral titres initially recovered during air sampling.

Here, we used the SARS-CoV-2 Delta (Pango B.1.617.2 [13]) as previous inhouse attempts to aerosolise and isolate SARS-CoV-2 Omicron (Pango BA1.1.529) had failed. As VoCs have different growth kinetics and plaque morphology it’s not clear if this is due to an inability to aerosolise and recover viable Omicron virus or if this method needs to be optimised for each VoC. It’s important that as SARS-CoV-2 continues to evolve, so does our understanding and response to these challenges. We recommend evaluation of all cell techniques for novel variants and hope this can act as a framework for optimisation.

Human aerosols generated through speech are not uniform, and laboratory generated aerosols are not able to replicate the full range of particle sizes produced. The BLAM used here typically generates particles with a diameter between 0.01-10μm and, in the model used here, does not replicate the composition of viral aerosols generated by human exhalations. In addition, human generated aerosols differ between individuals, in-particular in patients with acute disease [15]. It is important that this optimised method can be applied to naturally occurring aerosol generation through the establishment of a controlled cabinet where COVID-19 positive patients can talk and breath into whilst taking air samples.

It is hoped this optimised method for the recovery of laboratory generated SARS-CoV-2 aerosols can be used to support further research into SARS-CoV-2 transmission and inform methods of sampling within the environment.

